# Intramuscular prime/intranasal boost vaccination to induce sterilizing immunity against influenza A virus infection

**DOI:** 10.1101/2024.03.27.586965

**Authors:** Robin Avanthay, Obdulio Garcia-Nicolas, Nicolas Ruggli, Llorenç Grau Roma, Ester Párraga-Ros, Artur Summerfield, Gert Zimmer

## Abstract

The most commonly used influenza vaccines are made from inactivated viruses and are administered via the intramuscular route. Although these vaccines can protect from severe lower respiratory tract disease, they do not completely prevent virus replication in the upper respiratory tract, and this may lead to virus excretion and dissemination. Therefore, nasally administered live-attenuated influenza vaccines (LAIV) that induce mucosal immunity have been developed, but finding an optimal balance between sufficient attenuation and immunogenicity remained challenging. These problems apply to both human and swine influenza vaccines. We have recently developed an LAIV candidate based on the 2009 pandemic H1N1 virus which encodes a truncated NS1 protein and lacks PA-X protein expression (NS1(1-126)-ΔPAX). This virus showed a blunted replication and elicited a strong innate immune response. In the present study, we took advantage of the pig animal model to evaluate this vaccine candidate *in vivo* and to identify a strategy for its improvement. Nasal infection of pigs with the NS1(1-126)-ΔPAX LAIV candidate did not cause disease but was associated with prolonged virus shedding from the upper respiratory tract. To increase safety of the vaccine candidate, we developed a novel prime/boost vaccination strategy consisting of a haemagglutinin-encoding propagation-defective vesicular stomatitis virus replicon vaccine for primary immunization via the intramuscular route, and the NS1(1-126)-ΔPAX LAIV for secondary immunization via the nasal route. This immunization strategy significantly reduced LAIV shedding, increased the production of specific serum IgG, neutralizing antibodies, Th1 memory cells, and induced virus-specific mucosal IgG and IgA. Of particular note, the immune response induced by this vaccination strategy completely blocked replication of the homologous challenge virus in the respiratory tract, indicating that sterilizing immunity was achieved. In summary, our novel intramuscular prime/intranasal boost vaccine combines the features of high efficacy and safety which are urgently needed to combat influenza epidemics and pandemics.

**Author summary:** Inactivated influenza vaccines which are administered intramuscularly are safe but offer only limited protection. In addition, they do not adequately prevent virus transmission by infected individuals. On the other hand, nasally administered live-attenuated influenza vaccines induce a mucosal immune response, which can effectively prevent primary infection and virus excretion. However, live-attenuated vaccines might not be sufficiently immunogenic if they are too attenuated or they trigger a robust immune response but are still too virulent. To overcome this challenge, we have developed a novel prime/boost vaccination strategy consisting of an initial intramuscular immunization with a propagation-defective RNA virus vector and a subsequent nasal immunization with a modified influenza virus that has lost its ability to counteract the hosts‘ innate immune response. Using the pig model, we demonstrate that this approach elicited a more robust immune response both systemically and at mucosal surfaces. Importantly, replication of the vaccine virus in the respiratory tract was reduced, and challenge virus remained undetectable. In summary, our innovative vaccine, which combines intramuscular and intranasal routes of application, demonstrates high efficacy and safety and represents a valuable tool to control influenza epidemics and pandemics.

## Introduction

In humans, influenza A virus (IAV) causes acute respiratory infections that are normally associated with sudden high fever, muscle pain, headache, coughing, and fatigue (1–3). These symptoms begin from one to four days after exposure to the virus and may last for about 2–8 days. However, IAV can cause life-threatening infections, in particular if the virus disseminates to the lower respiratory tract which could lead to viral pneumonia and acute respiratory distress syndrome (ARDS) (4–6). Persons at high risk to suffer from severe influenza include the very young (< 2 years of age), the elderly (>65 years of age) or persons with underlying diseases such as diabetes, asthma, and cardiovascular diseases. IAV infections occur as seasonal epidemics which – in the Northern hemisphere – start in November and last until March. The cold season is associated with staying indoors, breathing dry air and having closer contacts, all factors that promote aerosol transmission which boost infection rates (7).

The most used human influenza vaccines to control seasonal epidemics represent inactivated viruses that are standardized according to their hemagglutinin (HA) content (8,9). These vaccines are administered to persons prior to the influenza season by intramuscular injection. The immunization triggers the production of serum antibodies that predominantly bind to the HA globular head domain and exhibit virus-neutralizing activity (10,11). A minor fraction of these virus-specific serum IgG is secreted into the lower respiratory tract where they provide protection from severe influenza pneumonia and ARDS (12,13). However, secretion of serum IgG into the mucosal tissues of the upper respiratory tract is not efficient (14–16).

Consequently, intramuscular immunization does not efficiently interfere with IAV replication in these tissues with the consequence that infectious virus is shed from the upper respiratory tract and potentially transmitted to other individuals. Not only those inactivated vaccines are unable to interrupt the infectious chain during seasonal epidemics, they may also drive antigenic drift by selecting for viral escape mutants typically with mutations in the HA globular head domain (17–19). To compete with the constant antigenic drift of HA, influenza vaccines must be updated every year. However, as the corresponding seed viruses are usually selected six months in advance, there is a considerable risk of antigenic mismatch with the circulating IAV strains (20,21). For some seasons, the efficacy of influenza vaccines can be relatively low, in particular in the elderly population (22,23).

In addition to inactivated vaccines, live-attenuated influenza vaccines (LAIV) have been approved (24,25). These vaccines are based on cold-adapted viruses which harbor temperature-sensitive mutations in the genes that encode the viral RNA polymerase complex. Following nasal administration, these cold-adapted LAIV only replicate in the upper respiratory tract without disseminating to the lung. The high level of attenuation of these vaccines limits LAIV transmission to contacts, but it also negatively impacts their immunogenicity (25). In addition, this vaccine is recommended for use in persons between 2 and 49 years of age. Thus, the elderly population is excluded from vaccination with these LAIV, although this population is particularly affected by severe influenza.

LAIV vaccines based on A/swine/Texas/4199-2/98 (H3N2) (sTX98) encoding a truncated NS1 protein were shown to protect pigs against challenge with the homologous virus (26,27) and to even provide protection against H1N2 virus (28–30). Such a vaccine was approved for use in piglets of one day of age in the US in 2017 (Ingelvac Provenza, Boehringer Ingelheim) but was withdrawn from the market following the discovery of reassortant viruses with gene segments of the vaccine and field strains (30,31). A recent study showed that NS1-truncated sTX98 replicated for a similar time than wild-type virus (32). These findings suggest that NS1-truncated LAIV are not sufficiently attenuated.

In our previous work, we generated recombinant LAIV candidates that were based on the pandemic A/Hamburg/4/2009 (H1N1) strain (pH1N1/09) (33). We compared a LAIV candidate encoding a C-terminally truncated NS1 protein (pH1N1/09-NS1(1-126)) with another pH1N1/09 mutant that had additional mutations in the PA gene that prevented PA-X expression (pH1N1/09-NS1(1-126)-ΔPAX). The latter virus further enhanced innate immune responses, replicated less efficiently, and induced less apoptotic cell death in a swine bronchiolar epithelial cell line. Based on these characteristics, we proposed that this virus could be a sufficiently attenuated LAIV candidate (33).

In the present study, these LAIVs were characterized and further developed using pigs as animal model. While both LAIV were highly immunogenetic, they appeared to be insufficiently attenuated with respect to the height and duration of vaccine virus shedding. We therefore developed a novel prime/boost vaccination protocol, which was based on a first intramuscular immunization with a propagation-defective vesicular stomatitis virus (VSV) vector encoding the pH1N1/09 HA antigen and a secondary immunization via the intranasal route using the LAIV candidates. The selection of the VSV vector was based on our previous work demonstrating efficacy in the induction of antibody and T cell responses to the HA antigen that mediated partial protection against heterologous IAV challenge of pigs in the absence of neutralizing antibodies (34). Our results identified a highly efficient vaccination protocol inducing strong systemic and mucosal immune responses, resulting in sterilizing immunity in pigs.

## Results

### Respiratory tract shedding of LAIV NS1(1-126)

In a recent work, we demonstrated that a recombinant pH1N1/09 encoding a modified NS1 protein that was truncated by 93 amino acids at the C-terminus replicated in a porcine bronchiolar cell line as efficiently as wild-type virus despite the induction of type I and III interferon (IFN-I and IFN-III) (33). To determine whether this LAIV candidate would be more attenuated *in vivo*, we infected pigs via the nasal route with either wild-type pH1N1/09 or the NS1(1-126) mutant. Monitoring of rectal temperature for 12 days revealed no fever in any of the animal groups (Fig 1A). Analysis of oropharyngeal swab samples by RT-qPCR showed that pH1N1/09 RNA was shed for up to 10 days post infection (Fig 1B). Compared to wild-type virus, the NS1(1-126) mutant was excreted at significantly lower levels with a reduction of viral RNA by almost two log_10_ between days 2 and 5, and a shortening of the overall excretion time by two days. Analysis of serum IgG by ELISA revealed that the NS1(1-126) mutant induced significantly lower levels of pH1N1/09-specific antibodies than wild-type virus (Fig 1C). Over time the serum IgG levels induced by the LAIV candidate dropped faster than those induced by wild-type virus. Nevertheless, in the bronchoalveolar lavage (BAL), the levels of virus-specific IgG were similarly high in both groups (Fig 1D). Likewise, virus-specific serum and BAL IgA levels did not differ significantly between the pH1N1/09- and NS1(1-126) mutant-infected animals (Fig 1E, F). We also measured the levels of neutralizing antibodies in serum and BAL fluids and found lower levels with the NS1(1-126) mutant when compared to pH1N1/09 group (Fig 1G, H). With respect to serum samples, the differences between the animal groups were only significant when the area under the curve (AUC) was analyzed (Fig 1G). Altogether, these data indicate that the LAIV candidate NS1(1-126) is still shed from the respiratory tract for a considerable time, albeit at reduced levels when compared to wild-type virus. Additionally, the systemic antibody response to the LAIV candidate was reduced when compared to that induced by wild-type virus infection.

**Figure 1.**
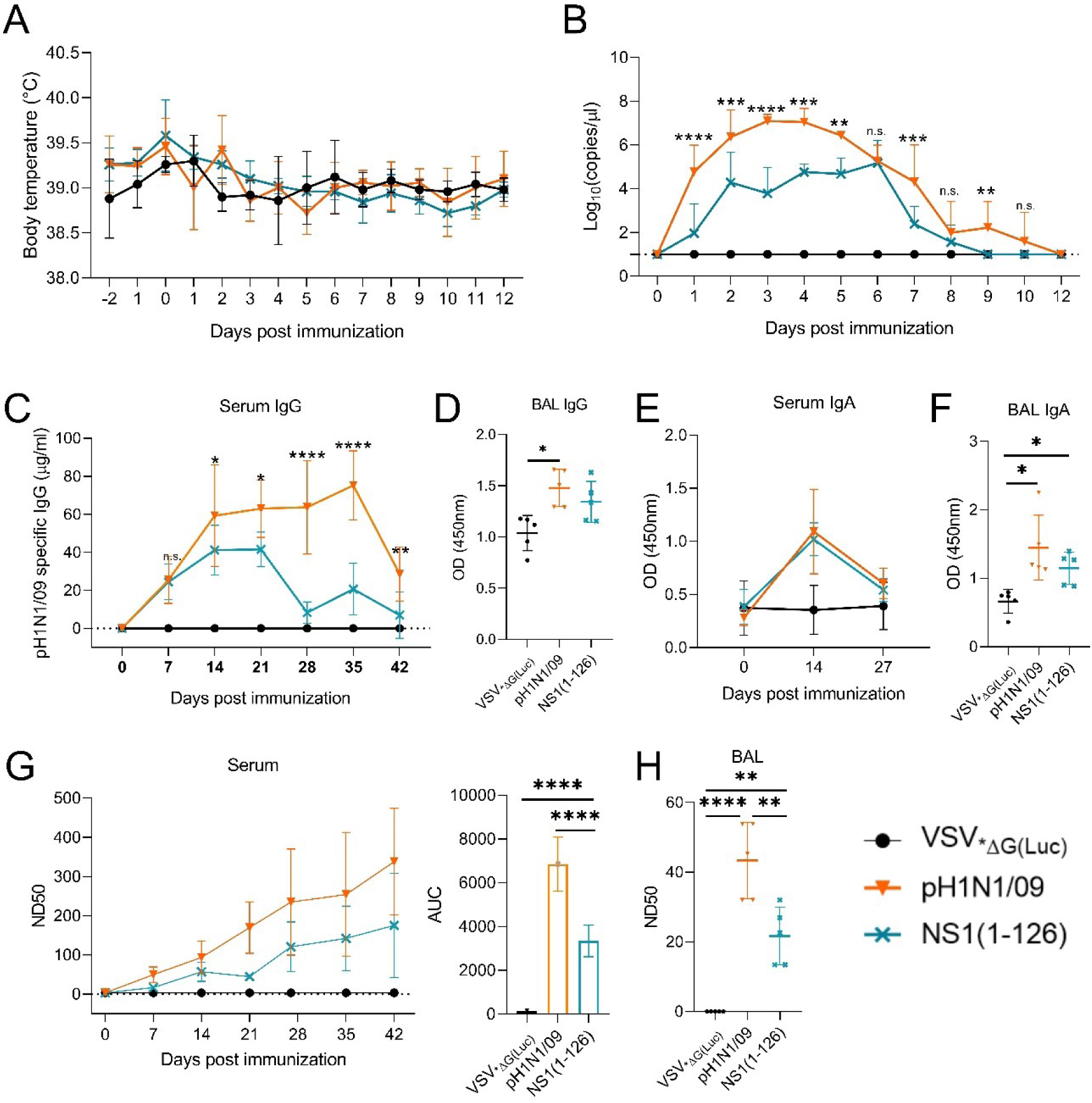
Replication and immunogenicity of pH1N1/09-NS1(1-126) in pigs following intranasal application. Nasal swabs samples were collected daily and serum samples weekly. At day 42 the animals were euthanized, and BAL fluid was collected. **(A)** Daily rectal body temperatures. **(B)** Viral RNA quantities in oropharyngeal swab samples. **(C, D)** pH1N1/09-specific Ig in sera and BAL fluids, respectively. **(E, F)** pH1N1/09-specific IgA in serum and in BAL fluids, respectively. **(G)** pH1N1/09-neutralizing antibody titres in the serum. The left panel depict average values at the individual days and the right panel the area under the curve (AUC) analyses encompassing all days. **(H)** pH1N1/09-neutralizing antibody titres in the BAL fluids. Statistical analyses employed two-way ANOVA tests for data in (B), (C) and (E); one-way ANOVA tests for (D), (F), (G, right panel) and (H). *p<0.05, **p<0.01, ***p<0.001, ****p<0.0001 indicate significant differences.

### Reduction of shedding following intramuscular prime/intranasal boost vaccination

To improve the performance of LAIV in terms of immunogenicity and reduced virus shedding, we developed a novel prime/boost immunization protocol (Fig 2A). The animals were first immunized via the intramuscular route with VSV_*ΔG(H1)_, a propagation-incompetent VSV vector encoding the pH1N1/09 HA in place of the VSV glycoprotein (G). Control animals were immunized with VSV_*ΔG(Luc)_ encoding firefly luciferase. Immunization of pigs with either VSV_*ΔG(H1)_ or VSV_*ΔG(Luc)_ did not cause an increase of body temperature (Fig 2B) or any other signs of disease. Four weeks after the primary vaccination, the pigs were immunized via the nasal route with the LAIV vaccine candidates (Table 1). The LAIV NS1(1-126) encoded a truncated NS1 protein lacking 93 amino acids at the C-terminus, and NS1(1-126)-ΔPAX combined both NS1 truncation and PA-X deletion (33). No fever was recorded for the first 10 days following infection with any of the LAIV (Fig 2C). Detection of viral RNA in oropharyngeal swab samples by RT-qPCR showed that animals which were first primed with the control vector VSV_*ΔG(Luc)_ and subsequently immunized via the nasal route with NS1(1-126)-ΔPAX shed virus at relatively high levels and for prolonged time (Fig 2D). In contrast, if the pigs were primed with VSV_*ΔG(H1)_ and then boosted with NS1(1-126)-ΔPAX, virus shedding was significantly reduced with no viral RNA detectable after day 6 (Fig 2E). Two out of five pigs showed virus shedding for only 1 day while one pig did not shed any virus (Fig 2E). Interestingly, if VSV_*ΔG(H1)-_primed animals were boosted with NS1(1-126), shedding was higher when compared to the VSV_*ΔG(H1)_/NS1(1-126)-ΔPAX vaccination protocol (Fig 2F). Together, these data demonstrate that shedding of LAIV can be efficiently reduced by an intramuscular prime/intranasal boost vaccination regimen as supported by analysis of their respective shedding (Fig 2G, area under curve).

**Figure 2.**
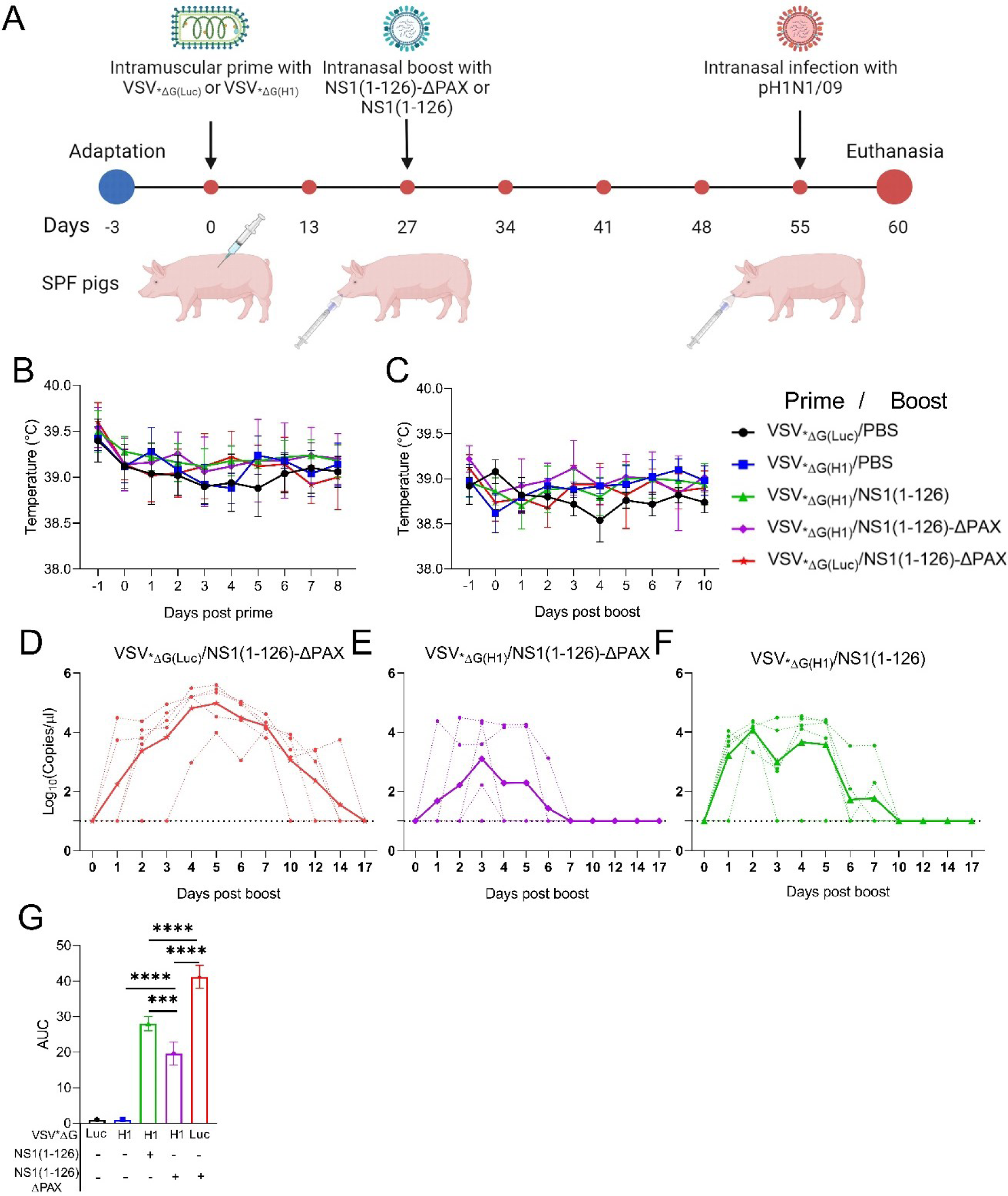
Intramuscular prime/intranasal boost vaccination protocol reduces LAIV shedding. **(A)** Schematic representation of the experiment design. Red points on the timeline indicate sampling days for blood (figure created on Biorender.com). **(B)** Body temperatures of pigs during the first eight days following primary immunization (i.m.). **(C)** Body temperature of pigs during the first 10 days after the subsequent intranasal LAIV immunization. **(D-F)** Viral RNA copies in oropharyngeal swab samples collected after the LAIV intranasal inoculation. Individual animals are represented by dashed lines and group mean values by continuous thick lines. In (D), the pigs were first immunized with negative control VSV_*ΔG(Luc)_ followed by pH1N1/09-NS1(1-126)-ΔPAX; in (E) with VSV_*ΔG(H1)_ followed by pH1N1/09-NS1(1-126)-ΔPAX, and in (F) with VSV_*ΔG(H1)_ followed by pH1N1/09-NS1(1-126). **(G)** AUC analyses for viral RNA quantities from days 0 to 17 after LAIV application (calculated with data from D-F). Significant differences of the AUC values were determined using the one-way ANOVA test (*p<0.05, **p<0.01, ***p<0.001, ****p<0.0001).

**Table 1.** Vaccine groups used in this study.

| Groups | Prime (name; abbreviations) | Boost (name; abbreviations) |
| --- | --- | --- |
| 1 | VSV_GFP_ΔG(Luciferase);<br>VSV*ΔG(Luc) | VSV_GFP_ΔG(Luciferase);<br>VSV*ΔG(Luc) |
| 2 | VSV_GFP_ΔG(H1);<br>VSV*ΔG(H1) | VSV_GFP_ΔG(Luciferase);<br>VSV*ΔG(Luc) |
| 3 | VSV_GFP_ΔG(H1);<br>VSV*ΔG(H1) | pH1N1/09-NS1(1-126);<br>NS1(1-126) |
| 4 | VSV_GFP_ΔG(H1);<br>VSV*ΔG(H1) | pH1N1/09-NS1(1-126)- ΔPAX;<br>NS1(1-126)-ΔPAX |
| 5 | VSV_GFP_ΔG(Luciferase);<br>VSV*ΔG(Luc) | pH1N1/09-NS1(1-126)- ΔPAX;<br>NS1(1-126)-ΔPAX |

### Enhanced systemic antibodies following intramuscular prime/intranasal boost vaccination

To assess the immune response following vaccination, serum and saliva were collected at different time points after primary and secondary immunization and analyzed by indirect ELISA. After the primary immunization with VSV_*ΔG(H1)_, virus-specific IgG were not detected (Fig 3A, B), but were strongly induced following intranasal boost with NS1(1-126)-ΔPAX or NS1(1-126). In contrast, significantly lower levels of specific serum IgG were detected if the animals were first immunized with the control vector VSV_*ΔG(Luc)_ and subsequently intranasally boosted with NS1(1-126)-ΔPAX (Fig 3A, right panel). With respect to serum IgA, the VSV_*ΔG(Luc)_/NS1(1-126)-ΔPAX vaccination protocol resulted in higher virus-specific IgA levels compared to the VSV_*ΔG(H1)_/NS1(1-126) and VSV_*ΔG(H1)_/NS1(1-126)-ΔPAX vaccination protocols (Fig 3B). No significant differences in the immune response following the boost immunization with NS1(1-126) or NS1(1-126)-ΔPAX LAIV were observed (Fig 3B). Interestingly, the serum IgA response was relatively short lasting with a peak at two weeks followed by a sharp decline. In contrast, serum IgG levels reached a plateau two weeks after the boost and thereafter decreased only slowly (Fig 3A).

**Figure 3.**
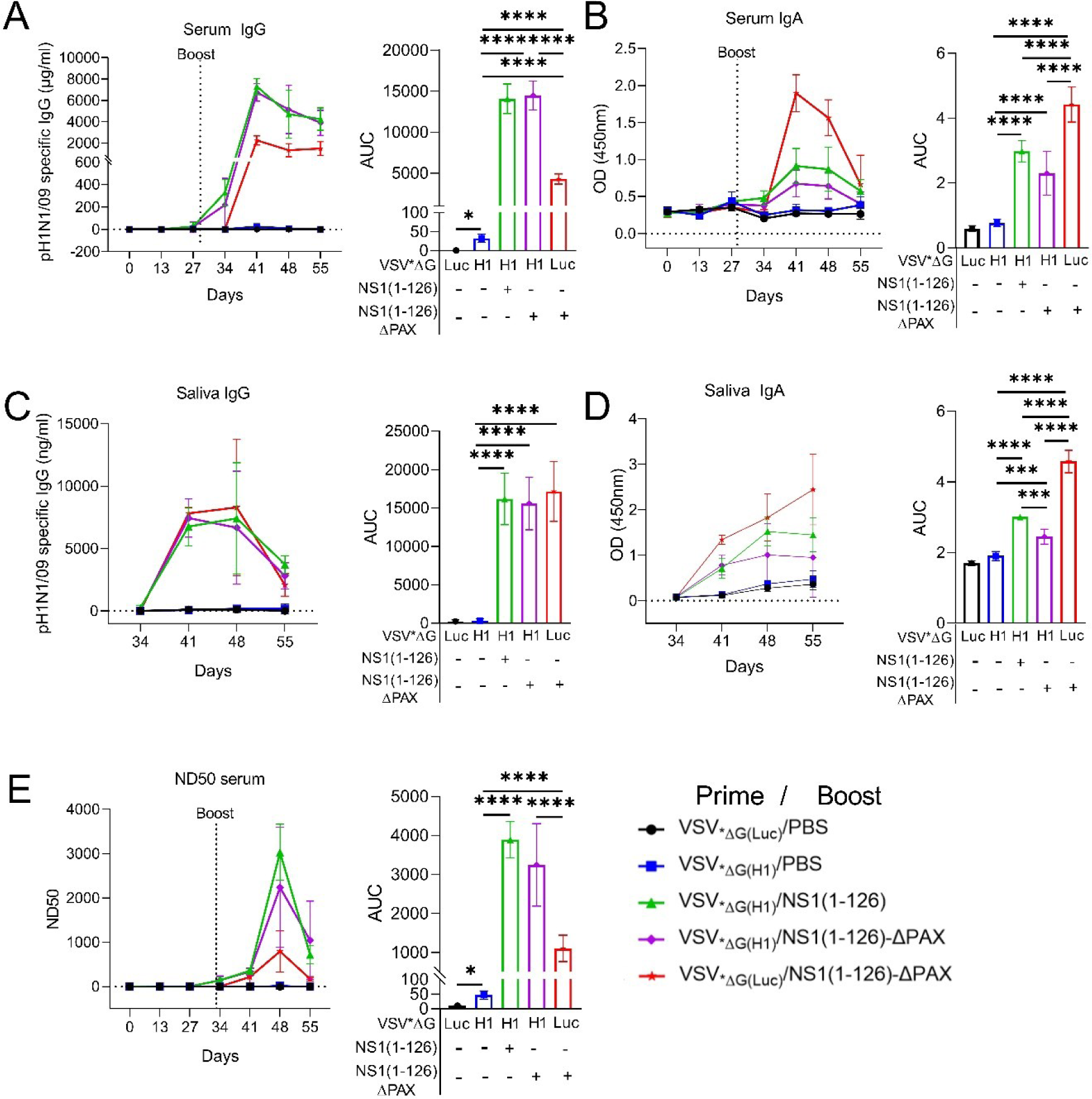
Antibody responses following intramuscular prime/intranasal boost vaccination. The experimental layout was as described in Fig 2A. In A-E the left panels show the kinetics of average antibody levels, whereas the right panels depict the AUC analyses of this data using all time points. **(A, B)** pH1N1/09-specific IgG or IgA in serum quantified by ELISA. **(C, D)** pH1N1/09-specific IgG or IgA in saliva quantified by ELISA. **(E)** Serum neutralizing antibody titres against homologous virus. Significant differences were determined using the one-way ANOVA test (*p<0.05, **p<0.01, ***p<0.001, ****p<0.0001).

In addition to serum antibodies, we also analyzed the mucosal immune response by measuring the pH1N1/09-specific IgG and IgA antibody levels in saliva by indirect ELISA. Intramuscular immunization with VSV_*ΔG(H1)_ followed by mock-vaccination with PBS via the nasal route did not result in the induction of detectable levels of IgG in saliva (Fig 3C). In contrast, when animals were primed with VSV_*ΔG(H1)_ and then boosted with either NS1(1-126) or NS1(1-126)-ΔPAX mutants, a strong increase in virus-specific secreted IgG was observed between days 34 and 41 post primary vaccination. These antibody levels stayed high until day 48 and then dropped (Fig 3C). No significant differences between the different LAIVs were observed. With respect to virus-specific IgA in saliva, an increase in ELISA reactivity was observed for the animals which were primed with either VSV_*ΔG(Luc)_ or VSV_*ΔG(H1)_ and boosted with PBS (Fig 3D). Considering that this reaction was also found in absence of H1 expression, it can be considered as a non-specific reaction. In contrast, the animals primed with VSV_*ΔG(H1)_ and subsequently boosted with either NS1(1-126) or NS1(1-126)-ΔPAX showed increased IgA responses. The highest IgA levels were induced by the VSV_*ΔG(Luc)_/NS1(1-126)-ΔPAX vaccination regimen (Fig 3D).

Next, we analyzed the induction of virus-neutralizing antibodies. Priming with VSV_*ΔG(H1)_ resulted in only very low levels of neutralizing antibodies (Fig 3E, right panel). However, if the H1-primed animals were boosted via the nasal route with either the NS1(1-126)-ΔPAX or the NS1(1-126) mutants, a significant increase in virus-neutralizing serum antibodies was detected. The ND50 values reached a peak of approximately 3000 at day 48 and dropped to an ND50 value of approximately 1000 at day 55 post primary immunization. In contrast, immunization with NS1(1-126)-ΔPAX of pigs that were injected with the control vector VSV_*ΔG(Luc)_ resulted in significantly lower neutralizing antibody levels compared to VSV_*ΔG(H1)_-primed and NS1(1-126)-ΔPAX-boosted animals.

### Enhanced CD4^+^ T-cell memory following intramuscular prime/intranasal boost vaccination

The cellular arm of the immune system is essential to help in the activation of the B cell through the production of cytokines and effector molecules (CD4^+^ T cell), and to clear virus-infected cells (CD8^+^ T cell) (35). To assess the level of CD4^+^ and CD8^+^ T-cell memory induction by the tested vaccinations, PBMCs were collected at day 48, restimulated with viral antigen and analyzed by flow cytometry (FCM) for intracellular expression of IFNγ, TNF and IL-17 in CD4^+^ or CD8^+^ T cells (Fig 4). Pigs vaccinated only once with VSV_*ΔG(H1)_ showed no significant activation of CD4^+^ and CD8^+^ T cells when compared to the control vector group (VSV_*ΔG(Luc)_). Animals that were primed with VSV_*ΔG(H1)_ and boosted with either NS1(1-126) or NS1(1-126)-ΔPAX LAIV showed significantly increased numbers of CD4^+^ T cells expressing TNF, IFNγ and both TNF and IFNγ but not IL-17s (Fig 4A-D). Importantly, this was not observed in the group that was primed with the control vector VSV_*ΔG(Luc)_ followed by immunization with NS1(1-126)-ΔPAX (Fig 4, A, B, D; red), demonstrating improved priming of CD4 Th1 cells by the intramuscular prime/intranasal boost protocol. No recall responses were observed for any of the vaccine groups in the CD8^+^ T cell subset (Fig 4E).

**Figure 4.**
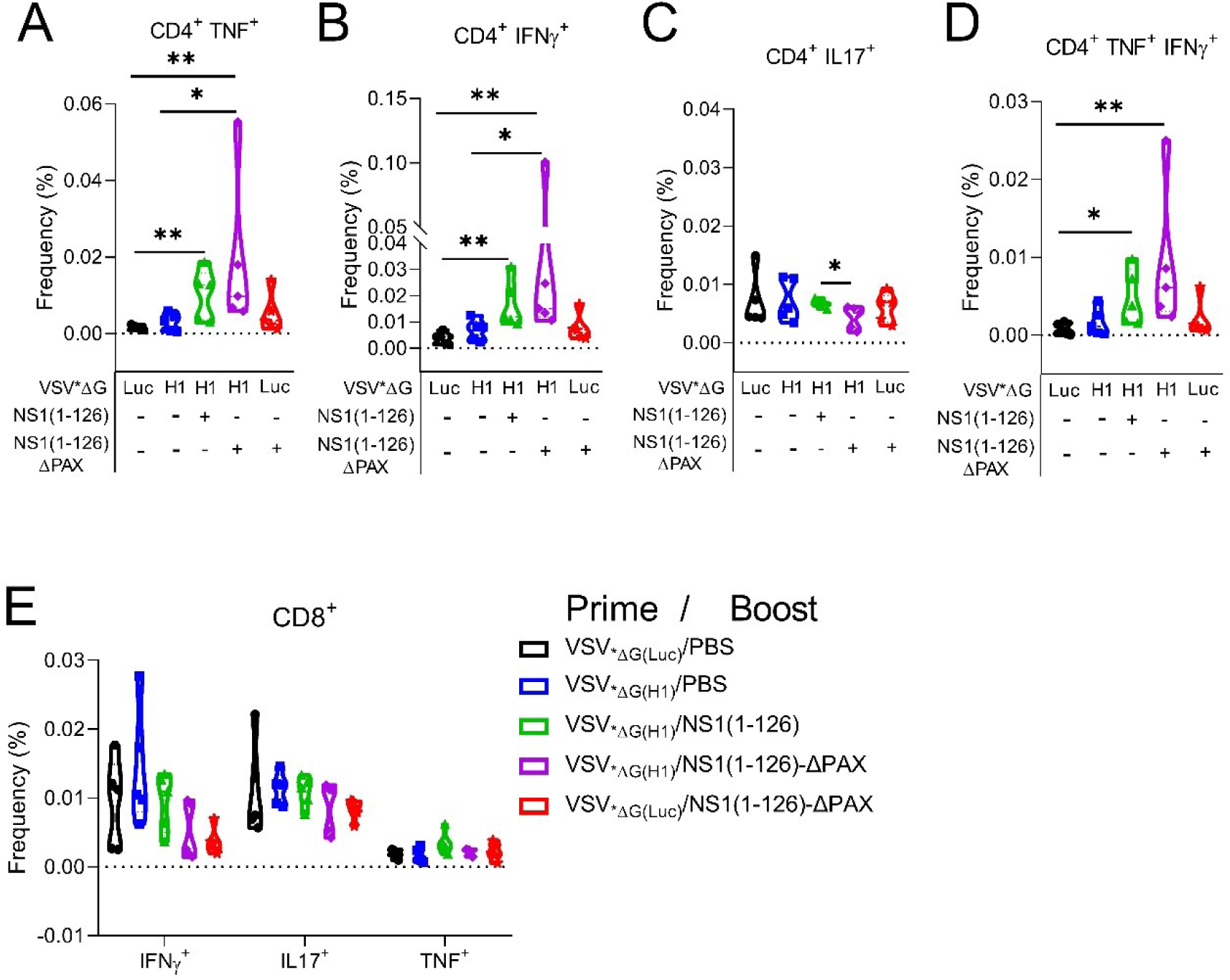
Induction of memory T-cell responses in the peripheral blood of pigs by intramuscular prime/intranasal boost vaccination. PBMCs were isolated from immunized pigs at day 48 and restimulated with live pH1N1/09 (see scheme in Fig 2A). **(A-D)** Intracellular cytokine staining of CD4^+^ T cells for TNF (A), IFNγ (B), IL17 (C), and TNF/IFNγ double-positive cells (D). The frequency of cytokine-positive cells relative to the total number of CD4^+^ T cells is shown. **(E)** Frequency of reactivated CD8^+^ cells that were positive for the indicated cytokines. Significant differences were determined using Mann-Whitney tests (*p<0.05, **p<0.01, ***p<0.001, ****p<0.0001).

### Induction of sterilizing immunity

To assess whether the intramuscular prime/intranasal boost vaccination would provide protection against infection with the homologous IAV strain, pigs were intranasally inoculated with pH1N1/09 using a dose of 10^6^ TCID_50_ per animal. Regardless of the treatments, none of the infected animals developed clinical signs of disease or fever (Fig 5A). For the vaccine groups VSV_*ΔG(H1)_/NS1(1-126)-ΔPAX, VSV_*ΔG(H1)_/NS1(1-126), and VSV_*ΔG(Luc)_/NS1(1-126)-ΔPAX, no viral RNA was detected in the swab samples (Fig 5B), indicating the absence of any virus replication in the respiratory tract of the immunized animals. In contrast, in the control group and the VSV_*ΔG(H1)_-primed only group, high level of virus shedding was observed which reached 10^6^ genome equivalents/µl at day two post infection and stayed at this level until euthanasia at day five (Fig 5B). When the VSV_*ΔG(H1)_-primed only group was compared to the control group a minor but significant reduction of viral RNA was found at days two and five post infection, and in the AUC analyses (Fig 5B). Following euthanasia of the pigs at day 5 post infection, viral RNA was also measured in lung homogenates. The highest levels were detected in animals of the control vaccine group, while the VSV_*ΔG(H1)_-primed only group displayed lower levels of viral RNA (Fig 5C). However, this difference was not significant due to high sample variability. Importantly, as observed in the swab samples, no viral RNA was detected in the lungs of animals immunized with all protocols utilizing LAIVs (Fig 5C). When viral RNA was measured in BAL fluids, the animals of the VSV_*ΔG(H1)_ -primed only group showed a significantly reduced level of viral RNA compared to the control group.

**Figure 5.**
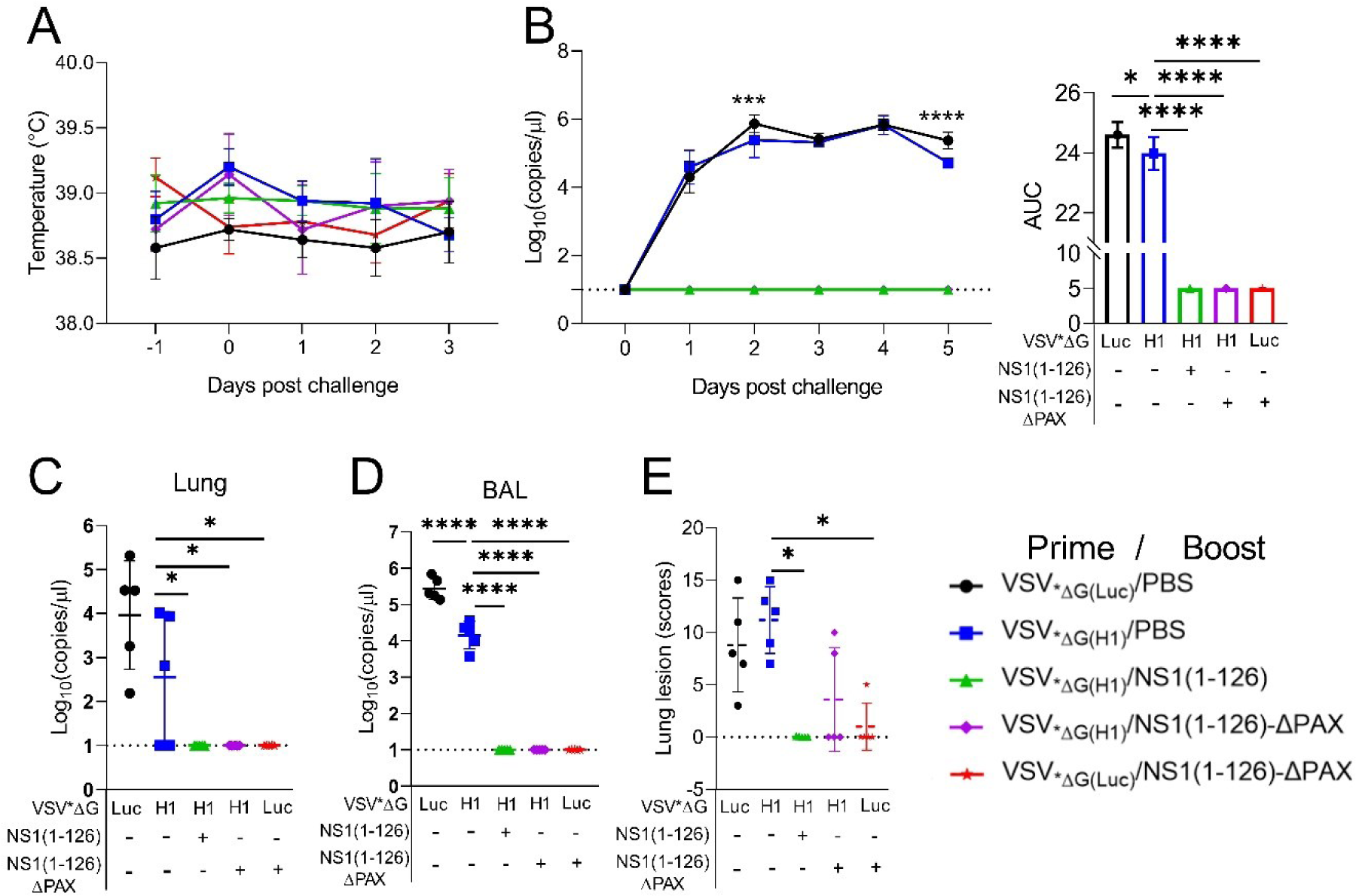
Efficacy of vaccine candidates following challenge infection. Pigs were vaccinated and immunologically monitored as described in Fig 2-4, and then challenged intranasally with pH1N1/09. **(A)** Body temperature after challenge infection. **(B)** Viral RNA loads in swab samples collected at all day’s post infection (left panel), and AUC analyses of this data (right panel). **(C, D)** Viral RNA loads in lung tissue and BAL fluids collected at day 5 post infection. **(E)** Histopathological lung lesion scores. Significant differences were determined using the one-way ANOVA test (AUC values in **B, C, and D**), and Kruskal-Wallis test (**E**) *p<0.05, **p<0.01, ***p<0.001, ****p<0.0001 indicate significant differences.

In contrast, as with the lung tissue, viral RNA was completely absent from the BAL fluids of all three vaccine groups that received the LAIV candidates (Fig 5D).

In agreement with the lack of clinical signs, the histopathological examination of the lungs identified only mild lesions typical of interstitial pneumonia. The histological lesions scoring was significantly higher in the VSV_*ΔG(H1)_-primed only than in LAIV groups (Fig 5E).

However, the macroscopic lesions, when present, were only mild and focal or multifocal. This may be a reason why there were no more differences between groups.

Postmortem analysis of mucosal antibody titers in BAL fluid revealed that pigs, which were first primed with VSV_*ΔG(H1)_ and then boosted with NS1(1-126)-ΔPAX LAIV, had significantly higher IgA levels compared to the VSV_*ΔG(H1)_ -prime only group (Fig 6A). In lung homogenates, only the VSV_*ΔG(Luc)_/NS1(1-126)-ΔPAX induced significantly higher IgA levels when compared to the negative control and the VSV_*ΔG(H1)_/PBS groups (Fig 6B). With respect to lung IgG levels, no significant enhancement was found even though animals that have been primed with VSV_*ΔG(H1)_ and boosted with either NS1(1-126) or NS1(1-126)-ΔPAX present an increased tendency (Fig 6C).

**Figure 6.**
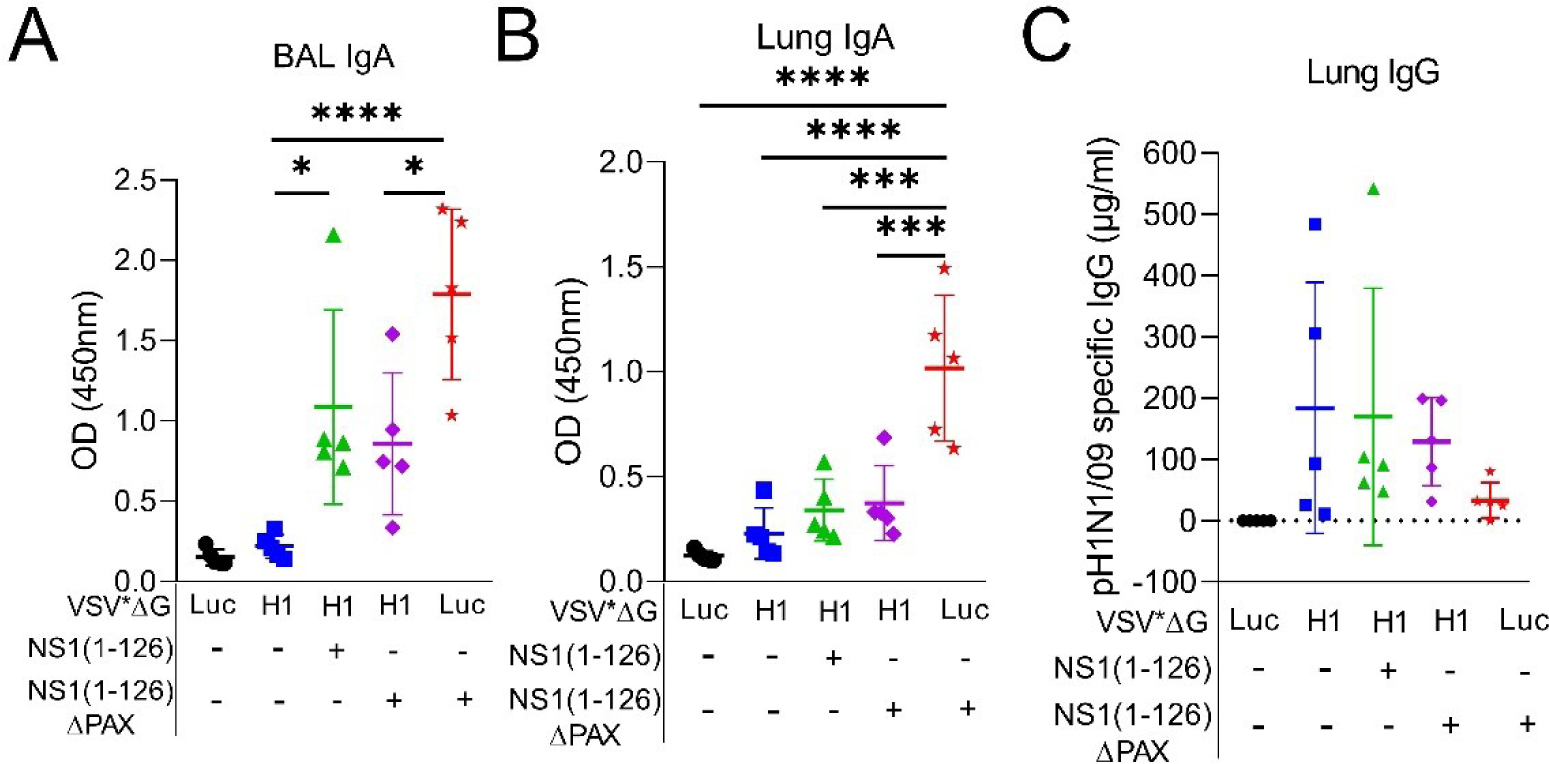
Antibody responses of pigs after nasal challenge infection with pH1N1/09. **(A, B)** pH1N1/09-specific IgA in BAL fluids and lung tissues 5 days post infection. **(C)** pH1N1/09-specific IgG in lung tissues. Significant differences were determined using the one-way ANOVA test. *p<0.05, **p<0.01, ***p<0.001, ****p<0.0001 indicate significant differences.

Taken together, these results demonstrate that our novel intramuscular prime/intranasal boost vaccination protocol induces enhanced systemic IAV-specific immunity in terms of higher IgG, higher neutralizing antibodies titers and enhanced memory Th1 cells. These systemic responses combined with the induction of local mucosal immunity in the respiratory tract provided sterilizing immunity against challenge infection. Importantly, both the priming with the VSV_*ΔG(H1)_ and the use of the NS1(1-126)-ΔPAX for the intranasal LAIV vaccine component enhanced the safety by strongly reducing LAIV shedding following vaccination and completely preventing shedding of challenge virus.

## Discussion

LAIV vaccines offer several advantages compared to inactivated IAV vaccines (36). First, LAIV infection results in the synthesis of all influenza virus antigens in their native conformation and thereby stimulate a broader antibody and T-cell response, that will improve protection against heterologous field viruses. Second, such vaccines also induce mucosal immunity when applied via the intranasal route and are expected to more efficiently stop transmission (37). Third, LAIV vaccines do not require adjuvants that can cause local and systemic side effects (38). Fourth, LAIV may have an improved duration of immunity due to prolonged antigenic stimulation and improved T-cell help which is required to generate long-lived plasma cells (39,40). On the negative side, LAIV vaccines can be problematic in terms of safety. If administered to patients with a weakened immune system, they might cause disease symptoms and be released from the upper respiratory tract and transmitted to other individuals (41). In addition, enhanced replication of LAIV may also favor the emergence of revertant virus that have regained virulence or may enhance the risk of formation of reassortant viruses in the case of co-infection with field virus. On the other hand, if replication of LAIV is too restricted due to over-attenuation, virus antigen levels will be too low to trigger a strong immune response. Therefore, finding the balance between safety and efficacy remains challenging.

An example for LAIV failure is the NS1-truncated LAIV pig vaccine that had to be withdrawn from the market after emergence a reassortant virus with gene segments from field IAV and the vaccine strains (30). In support of this, the present study shows that the shedding of this virus (NS1(1-126)) was lower than pH1N1/09 but still detectable for at least 8 days, and similar observations have been made by others using H3N2 swine IAV (32). All this demonstrates the importance of novel or additional attenuation strategies.

We tested one such approach by modifying the PA gene to eliminate expression of the accessory PA-X protein in the context of the pH1N1/09 NS1(1-126) mutant. Indeed, this NS1(1-126)-ΔPAX mutant stimulated higher levels of IFN and showed restricted replication in porcine bronchiolar epithelial cells (33). However, to our surprise, our present data demonstrated that NS1(1-126)-ΔPAX infected pigs still shed high levels of virus for prolonged time from the respiratory tract. A possible explanation could be that virus detected in swabs originated from the upper respiratory tract where low temperature may cause weak innate immune responses (42). Considering that a main attenuation target of the NS1(1-126)-ΔPAX mutant is the viral silencing of innate immunity, this might explain why the attenuation observed *in vitro* might not well translate to the *in vivo* situation, at least with respect to shedding.

When evaluating further strategies to develop a vaccine that fulfils the four points mentioned at the beginning of this Discussion, we were concerned about the observation that the NS1(1-126) mutant induced clearly weaker systemic and mucosal neutralizing antibodies when compared to wild-type virus. To avoid a further reduction of immunogenicity by additional attenuation, we developed a different strategy that was based on a heterologous prime/boost protocol. Our concept combines the high capacity of VSV-based vectors to induce systemic antibody and T cell responses after intramuscular injection (34) with the LAIV approach. To this end, we employed the highly safe propagation-defective VSV_*ΔG(H1)_ replicon particles expressing HA for intramuscular priming and NS1- and/or PA-X-modified LAIV for nasal boosting to induce local protection. This approach strongly reduced LAIV shedding in particular when the double mutant NS1(1-126)-ΔPAX was employed. As virus shedding was monitored by RT-qPCR analysis of swab samples, we cannot exclude that excreted virus was still infectious. Future experiments will show whether transmission of NS1(1-126)-ΔPAX from H1-primed to naïve animals takes place. In addition to reduced shedding, the heterologous VSV/LAIV vaccination protocol was more immunogenic in terms of induction of systemic IgG and neutralizing antibodies as well as memory Th1 cells. While mucosal antibody levels were below those induced by LAIV alone, the immunity induced resulted in complete protection from challenge infection as indicated by the absence of lung lesions and the lack of viral RNA in tissues of the respiratory tract. The achievement of such a sterilizing immunity would stop transmission chains and therefore have a major impact on slowing down epidemics.

Future studies are required to determine if the present prime/boost vaccination protocol may also have advantages to protect against antigen-drifted viruses. Even if sterilizing immunity to such viruses may not be achieved, we anticipate that compared to the immunization with current inactivated vaccines, shedding and transmission of IAV challenge viruses will be largely reduced. Thus, this novel prime/boost vaccination protocol will help to control seasonal influenza epidemics or pandemics more efficiently both in terms of improved effectiveness at preventing disease and better efficacy at the population level.

## Material and methods

### Ethics statement

The study was performed according to Swiss laws (the Animal Welfare Act TSchG SR 455, the Animal Welfare Ordinance TSchV SR 455.1, and the Animal Experimentation Ordinance TVV SR 455.163). All experiments were reviewed by the committee on animal experiments of the canton of Bern and approved by the cantonal veterinary authority under the license number BE128/19.

### Cells

Madin-Darby canine kidney type II cells (MDCK-II) were kindly provided by Georg Herrler (University of Veterinary Medicine, Hannover, Germany) and maintained with minimum essential medium (MEM, Thermo Fisher Scientific, Basel, Switzerland; cat. no. 31095-029) supplemented with 5% of fetal bovine serum (FBS; Pan Biotech, Aidenbach, Germany; cat. no. P30-3033). Human embryonic kidney (HEK) 293T cells (American Type Culture Collection (ATCC), Manassas, USA; cat. no. CRL-3216) were maintained with Dulbecco’s Modified Eagle Medium (DMEM, Thermo Fisher Scientific; cat. no. 32430-027) supplemented with 10% FBS. Baby hamster kidney 21 (BHK-21) fibroblasts were obtained from ATCC (cat. no. CCL-10) and maintained in Glasgow’s minimal essential medium (GMEM, Thermo Fisher Scientific, cat. no. 21710-025) supplemented with 5% FBS. BHK-G43 cells, a transgenic BHK-21 cell clone expressing VSV glycoprotein G in a regulated manner, was maintained in GMEM supplemented with 5% FBS (43). All cells were grown at 37°C in a 5% CO_2_ atmosphere.

### Generation of recombinant influenza virus

The pHW2000 plasmids encoding the 8 segments of A/Hamburg/4/2009 (H1N1) (GenBank accession nos.: GQ166207, GQ166209, GQ166211, GQ166213, GQ166215, GQ166217, GQ166219, GQ166221) were originally provided by Hans-Dieter Klenk (University of Marburg, Marburg, Germany). The pHW2000 plasmids encoding RNA segments 3 (PA) and 8 (NS1, NEP) were modified as previously described (33). Briefly, four stop codon were inserted at position 126 of the NS1 open reading frame leading to a truncated protein lacking the last 93 amino acids. This virus is termed “NS1(1-126)”. To eliminate PA-X expression, four nucleotides were changed at position 191 without modifying the amino acid sequence of the PA protein. This virus was termed “NS1(1-126)-ΔPAX”. Recombinant viruses were generated using the eight-plasmid system as previously described (33,44) and passaged two times on MDCK-II cells with FBS-deficient medium containing 1% penicillin/streptomycin and 1 µg/ml of N-tosyl-L-phenylalanine chloromethyl ketone-treated trypsin (TPCK-trypsin, Merck KGgA, Darmstadt, Germany, cat. no. 4370285). Infectious virus titers were determined on MDCK-II cells as previously described (45), using a monoclonal antibody directed to the influenza virus nucleoprotein clone (H16-L10-4R5; ATCC, HB-65) for detection of infected cells by indirect immunofluorescence.

### Generation of recombinant VSV vector vaccine

The HA gene of A/Hamburg/4/2009 (H1N1) (GenBank acc. no. GQ166213) was amplified by PCR using the Phusion DNA polymerase and inserted into the *Mlu*I and *Bst*EII sites of the pVSV*ΔG(HA_H5-HP_) plasmid (46), resulting in the pVSV*ΔG(H1) plasmid. Recombinant vesicular stomatitis virus (VSV) replicon particles were generated as previously described (34,47,48). Briefly, BHK-G43 cells were first infected with modified vaccinia virus Ankara encoding the T7 phage RNA polymerase (MVA-T7, kindly provided by Gerd Sutter from Ludwig-Maximilians-Universität, München, Germany) using a multiplicity of infection (moi) of 3 focus-forming units (ffu) per cell. Subsequently, the cells were transfected with pVSV*ΔG(H1) along with three plasmids encoding the VSV N, P, and L proteins; all under the control of the T7 promoter. The cells were incubated for 24h at 37°C and 5% CO_2_ in the presence of 10^-9^ M of mifepristone (Merck KGaA, Darmstadt, Germany) to allow expression of the G protein. Then, the cells were trypsinized and seeded along with fresh BHK-G43 cells into a T75 flask and incubated with GMEM containing 5% FBS and 10^-9^ M mifepristone for 24h at 37°C and 5% CO_2_. The supernatant was harvested, and cells debris removed by low-speed centrifugation, and the supernatant passed through a 0.2 µm pore-size filter. The recombinant VSV*ΔG(H1) (VSV_*ΔG(H1)_) vector was propagated on mifepristone-treated BHK-G43 cells and stored in aliquots at −70°C. Infectious virus titers were determined on BHK-21 cells. The recombinant VSV*ΔG(Luc) (VSV_*ΔG(Luc)_) vector has been generated previously and was propagated accordingly (47).

### Animal experiments

For the first animal experiment, 15 healthy 10-weeks old large white conventional pigs were purchased from Agroscope (competence center of the Swiss confederation in the field of agricultural and agri-food research). The animals were randomly allocated into three groups each containing five pigs of mixed sex. The animals were intranasally immunized with 10^6^ ffu/animal of either pH1N1/09, pH1N1/09-NS1(1-126), or VSV_*ΔG(Luc)_ as control. Body temperature and clinical symptoms of disease were monitored for the following 12 days. Oro-nasal swab samples were taken daily for the first 12 days post immunization to monitor virus shedding. Serum and saliva samples were taken once a week for a total period of six weeks after immunization to monitor for systemic and local antibody responses. Six weeks after immunization, the animals were euthanized by electroshock and subsequent exsanguination. Bronchoalveolar lavage (BAL) was prepared immediately after exsanguination.

For the second animal experiment, 25 healthy 10-weeks old specific pathogen-free (SPF) Swiss large white pigs from the Institut of Virology and Immunology (IVI, Mittelhäusern) breeding facility were used. The animals were tested seronegative for the influenza A virus nucleoprotein by using a commercial ELISA (ID Screen® Influenza A Antibody Competition Multi-species ELISA, ID-Vet, Montpellier, France, cat. no, FLUACA). The animals were divided randomly into five groups each containing five pigs of mixed sex and immunized via the intramuscular route using a dose of 10^8^ ffu/pig of either VSV_*ΔG(H1)_ or the control vector VSV_*ΔG(Luc)_ (**Table 1**). Temperature and clinical symptoms were monitored daily for seven days, and serum samples were prepared at days 13 and 27 after primary immunization (**Fig 2A**). At day 28 after primary immunization, the animals were immunized via the nasal route with LAIV (see **Table 1**) using a dose of 10^5^ ffu/pig.

To this end, an MAD Nasal^TM^ intranasal mucosal atomization device (Teleflex Medical Europe Ltd., Ireland, cat. no, MAD300) plugged to a 10-ml syringe was employed. Body temperature and clinical symptoms were monitored for one week following the second (boost) immunization. Swabs samples were taken daily, suspended in 2 ml of MEM, and stored at −70°C prior to use. Serum and saliva samples were taken every week for four weeks and stored at −20°C prior to serological testing. Blood samples were collected at day 49 (three weeks post boost) for isolation of PBMCs and T cell reactivation experiments. At day 56, the animals were infected with wild-type pH1N1/09 via the nasal route using 10^6^ ffu/pig. Body temperature and clinical symptoms were monitored, and serum and saliva samples were collected daily. Five days after challenge infection, the animals were euthanized by electroshock and subsequent exsanguination. Immediately after exsanguination, the lung’s right cranial lobe was prepared for viral RNA detection and lung samples were taken in formalin as indicated below for histological analysis. Bronchoalveolar lavage was prepared for detection of virus-specific antibodies and viral RNA.

### ELISA

For detection of antibodies directed against pH1N1/09, 96-well plates (Nunc^TM^ MaxiSorp^TM^) were coated overnight at room temperature with 1 µg/well of heat-inactivated A/H1N1/09 (56°C, 30 min). Plates were blocked for 1h at 37°C with 200 µl/well of 1% bovine serum albumin (BSA) in PBS containing 0.05% (v/v) Tween 20. Serum, saliva, and BAL samples were added in duplicates at the right dilution and incubated for 1h at 37°C. Ten-fold serial dilutions of pH1N1/09 HA-specific antibody (Thermo Fisher, cat. no. PA5-81645) were used as standard. Plates were washed with PBS (without Ca^2+^ and Mg^2+^) containing 0.05% (v/v) of Tween 20 and incubated with horse radish peroxidase (HRP)-conjugated anti-swine IgG (Abcam, cat. no. ab6915) or anti-swine IgA (Bethyl, cat. no. A100-102P) diluted 1/2000 and 1/50’000, respectively. The cells were washed three times with 200 µl/well of PBS/Tween and incubated for 10 min at room temperature with 50 µl/well of 3,3‘,5‘,5‘-tetramethylbenzidine (TMB)/H_2_O_2_ peroxidase substrate (Merck KGaA, cat. no. T4444). The reaction was stopped by addition of 50 µl/well of 1 M HCl and absorbance read at 450 nm using a GloMax microplate reader (Promega, Dübendorf, Switzerland).

### Virus neutralization test

Porcine sera were serially diluted in MEM medium (two-fold or five-fold dilution steps) and added in quadruplicates to 96-well microtiter plates (50 µl/well). To each well 50 µl of pH1N1/09 (2000 ffu/ml) were added and incubated for 1h at 37°C. The antibody/virus mix was then added to MDCK-II cells that were grown to confluence in 96-well cell culture plates and incubated for 1h at 37°C and 5% CO_2_. The cells were washed once and incubated with fresh medium (100 µl/well) for 24h at 37°C and 5% CO_2_. The cells were fixed with 4% formaldehyde solution and immuno-stained as previously described (45). The 50% neutralizing dose (ND_50_) was calculated using the Spearman-Kärber method (49).

### RNA extraction and RT-qPCR

All samples were directly transferred to 700 µl of RA1 lysis buffer (Macherey-Nagel, Düren, Germany, cat. no. 740961) containing 1% β-mercaptoethanol. Organs were homogenized in RA1 lysis buffer using a tissue bullet blender (Next Advanced Inc., Troy, NY, USA). Total RNA was extracted from the lysates using the NucleoMag Vet kit (Macherey-Nagel, cat.no. 744200) according to the manufacturer’s protocol. Reverse transcription from RNA to cDNA and real-time quantitative PCR (qPCR) were performed on a QuantStudio 5 real-time PCR system (Thermo Fisher Scientific) using the AgPath-ID™ One-Step RT-PCR kit (Life Technologies, Zug, Switzerland, cat. no. AM1005) with vRNA segment 7-specific oligonucleotide primers and probe (50,51). Data were acquired and analyzed using the Design and Analysis Software v1.5.2 (Thermo Fisher Scientific). Quantification was performed using an internal standard based on the IAV RNA segment 7.

### T cell assay

Peripheral blood mononuclear cells (PBMC) were isolated from 50 ml of blood in EDTA per pig. For each animal, five million PBMC were seeded per well (12-well plate format) in triplicates and stimulated for 14h at 39°C and 5% CO_2_ with pH1N1/09 (moi of 0.1 ffu/cell). Thereafter, brefeldin A at 3µg/ml (Invitrogen, cat. no. 00-4506-51) was added to the cells and incubated for another 4 h. The cells were washed twice and stained with the fixable Aqua Dead Cell Stain kit (ThermoFisher, cat. no. L34957). The CD4^+^ and CD8^+^ T lymphocyte cells were labeled by incubation with anti-CD4 IgG2b (clone 74-12-4, hybridoma kindly obtained from Dr. Joan Lunney, USDA Beltsville, MD, USA; (52)) and anti-CD8β IgG2a (PG164A, BioTechne, Basel, cat. no. NBP2-60955), followed by isotype-specific AlexaFluor-488 (Thermo Fisher, cat. no. A21141) and PE-Cy7 conjugates (Abcam, cat.no. ab130787). After fixation and permeabilization, anti-IFNγ-PE diluted 1/200 (P2G10, BD Biosciences, cat. no. 561481), anti-TNF-AF647 diluted 1/20 (Mab11, Biolegend, cat. no. 502916) and anti-IL17 diluted 1/5 (SCPL1362, BD Bioscience, cat. no. 560436) were added. Data were acquired on FACS Canto-II (BD Bioscience) and analyzed using FlowJo software, version 10 (BD Bioscience).

### Histopathology

Three lung samples of approximately 1 cm in diameter from the same location of the right lung: apical lobe, medial lobe, and caudo-dorsal area of the diaphragmatic lobe, were systematically taken and placed in 4 % formalin. In addition, when macroscopic lesions compatible with broncho interstitial pneumonia were observed, up to three additional samples from the affected lobes were taken for histological examination. The samples were trimmed and placed in two different cassettes per pig separately, with one cassette containing the three systematically collected lung sections and the other containing up to three lung sections with macroscopic lesions. The microscopic evaluation was made based on the previously reported Morgan score (53,54). Necrosis of the bronchiolar epithelium, airway inflammation, perivascular/bronchiolar cuffing, alveolar exudates, and septal inflammation, were scored from 0-4 points each (0: none, 1: minimal, 2: mild, 3: moderate, and 4: severe). All the parameters were added to obtain the scoring per slide (0-20 points), per animal (sum of slide 1 and 2, 0-40 points) and per group (mean score per animal for each group, 0-40 points).

### Statistical analysis

Data analysis and figures were done using GraphPad Prism 8 Software (GraphPad Software). One-way and two-way ANOVA tests with multiple comparisons, Kruskal-Wallis test, and Mann-Whitney tests were used to determine statistical significance in experimental data (detection of viral RNA by RT-qPCR, antibody titers, and T cell recall responses). P values lower than 0.05 was considered as statistically significant.

## Author contributions

RA acquired and analysed data and wrote the manuscript. GZ and AS conceived the idea, designed the study, and reviewed the manuscript. OG and NG proceed to the sampling of the animals and reviewed the manuscript. LG and EP performed the histopathological analysis. All authors gave their final approval for submission.

## Acknowledgments

We would like to thank Martin Schwemmle (University of Freiburg, Germany) for providing plasmids. We are grateful to Sylvie Python, Noelle Donze, Caroline Lehmann for their technical support.

